# Boosting heterologous protein production yield by adjusting global nitrogen and carbon metabolic regulatory networks in *Bacillus subtilis*

**DOI:** 10.1101/319293

**Authors:** Haojie Cao, Julio Villatoro-Hernandez, Ruud Detert Oude Weme, Elrike Frenzel, Oscar P. Kuipers

## Abstract

*Bacillus subtilis* is extensively applied as a microorganism for the high-level production of heterologous proteins. Traditional strategies for increasing the productivity of this microbial cell factory generally focused on the targeted modification of rate-limiting components or steps. However, the longstanding problems of limited productivity of the expression host, metabolic burden and non-optimal nutrient intake, have not yet been solved to achieve production strain improvements. To tackle this problem, we systematically rewired the regulatory networks of the global nitrogen and carbon metabolism by random mutagenesis of the pleiotropic transcriptional regulators CodY and CcpA, to allow for optimal nutrient intake, translating into significantly higher heterologous protein production yields. Using a β-galactosidase expression and screening system and consecutive rounds of mutagenesis, we identified mutant variants of both CcpA and CodY that in conjunction increased production levels up to 290%. RNA-Seq and electrophoretic gel mobility shift analyses showed that amino acid substitutions within the DNA-binding domains altered the overall binding specificity and regulatory activity of the two transcription factors. Consequently, fine-tuning of the central metabolic pathways allowed for enhanced protein production levels. The improved cell factory capacity was further demonstrated by the successfully increased overexpression of GFP, xylanase and a peptidase in the double mutant strain.

**Highlights:**

- The global transcription machinery engineering (gTME) technique was applied to build mutational libraries of the pleiotropic regulators CodY and CcpA in *Bacillus subtilis*
- Specific point mutations within the DNA-binding domains of CodY and CcpA elicited alterations of the binding specificity and regulatory activity
- Changes in the transcriptome evoked the reprogramming of networks that gear the carbon and nitrogen metabolism
- The rewired metabolic networks provided a higher building block capacity for heterologous protein production by adjusting the nutrient uptake and channeling its utilization for protein overexpression

## 1. Introduction

*Bacillus subtilis* is a well-characterized microbial cell factory that is widely used for the production of a variety of proteins for commercial and medical applications (Song et al., 2015; van Dijl and Hecker, 2013; Westers et al., 2004; Zweers et al., 2008). Improving the production potential of this classic chassis strain has been a research focus for several decades. Numerous engineering and biotechnological approaches have been employed in attempts to enhance production yields in industrial strains, for instance by utilizing modified promoters and RBSs, codon-optimization, pathway rerouting or gene disruption (Chen et al., 2015; Kang et al., 2014; Liu et al., 2017). However, although remarkable progress has been made in improving the protein overproduction capacity of *B. subtilis*, the space for traditional techniques or strategies to further improve this host organism’s productivity is increasingly limited.

In nature, the intracellular distribution of various resources in healthy cells has been ‘optimized’ by natural evolution over very long periods of time (Wu et al., 2016). Introducing an overexpression pathway for heterologous proteins into an engineered organism requires a large proportion of the host cell’s resources, including ATP, carbohydrates and amino acids. This imposed metabolic drain has been defined as ‘metabolic burden’ or ‘metabolic load’ (Zou et al., 2017). In this case, the vast majority of the intracellular metabolic fluxes, including energy resources such as NAD(P)H and ATP and carbon/nitrogen/oxygen building blocks, are forcibly assigned towards the heterologous product biosynthesis (Glick, 1995). The essential requirements for cellular maintenance, in turn, become imbalanced and insufficient in the engineered microbes (Pitera et al., 2007). Therefore, the biosynthetic yield of the expressed target product will remain at a relatively low level (Colletti et al., 2011; Glick, 1995), or even suddenly drop into the ‘death valley’ (minimal production level) on a ‘cliff’ under suboptimal growth conditions (Wu et al., 2016). Hence, the strategy to reduce the metabolic burden in a microbial host by enhancing the uptake of required nutrients and balancing heterologous and native metabolic flux demands, which could potentially benefit the robust production of large quantities of the target product.

In *B. subtilis*, the molecular mechanisms of nutrient-sensing based central metabolic regulations have become increasingly clear. The global transcriptional regulator CodY either represses or, less frequently, induces the transcription of target genes in the late exponential or early stationary phase in the presence of high intracellular levels of GTP and branched-chain amino acids (BCAAs; isoleucine, valine, and leucine) (Brinsmade et al., 2014; Shivers and Sonenshein, 2004). BCAAs act as corepressors by sterically triggering conformational changes that lead to altered DNA binding capabilities (Levdikov et al., 2017). This transcriptional regulation enables cells to adapt to various nutrient conditions in different growth environments, inducing a wide variety of cellular processes such as sporulation, competence development, nitrogen metabolism and biofilm formation (Belitsky and Sonenshein, 2013; Sonenshein, 2007). A second global transcriptional regulator that orchestrates fluxes in the central metabolism, specifically carbon utilization, is the extensively studied catabolite control protein A (CcpA). This transcription factor becomes active when in complex with phosphorylated histidine-containing protein (HPr) or its paralogous protein Crh (Mijakovic et al., 2002; Schumacher et al., 2004). This activity is enhanced by fructose-1,6-bisphosphate (FBP) and glucose-6-phosphate (G6P) when the cells are grown with glucose or other preferentially utilized carbon sources (Schumacher et al., 2007). Subsequently, activated CcpA binds to the cis-acting DNA-binding sites termed catabolite repression elements (*cre* sites) of the target regulon, leading to carbon catabolite repression (CCR) or carbon catabolite activation (CCA) (Fujita, 2009; Marciniak et al., 2012; Stulke and Hillen, 2000; Weme et al., 2015b). Both CodY and CcpA behave either as a repressor or activator of gene expression by specifically binding to a sequence located in or near the promoter region of target genes. Thus, these two global regulatory proteins and their ligands FBP, GTP and BCAAs, jointly control the intersections of large regulons that balance the use of available nutrient sources, systemically coordinate the intracellular carbon and nitrogen fluxes and contribute to cell homeostasis by stimulating specific catabolic processes.

Prior studies showed that global transcription machinery engineering (gTME) elicits a global alteration at the transcriptional level that perturbs the expression of multiple proteins simultaneously, which allows acquisition and selection of phenotypes of interest from a broad library (Alper and Stephanopoulos, 2007; Tyo et al., 2007). Some global transcription machinery components, such as sigma factors in bacteria (Alper and Stephanopoulos, 2007; Klein-Marcuschamer and Stephanopoulos, 2008), zinc finger-containing artificial transcription factors (Park et al., 2003), and Spt15 in yeast (Alper et al., 2006) were randomly mutagenized for generating phenotypes of biotechnological interest, including improved production capacities and strain tolerance towards elevated end-product levels. We thus hypothesized this strategy could be exploited to rewire the nitrogen and carbon metabolic flux distributions and to optimize nutrient uptake and utilization in *B. subtilis* at the whole-cell level to gain enhanced protein production traits by specific adjustments of the activity of CodY and CcpA. In addition, this study provides deeper insights into the interaction between CodY and CcpA by the analysis of the globally rewired nitrogen and carbon metabolic networks.

## 2. Materials and methods

### 2.1. Bacterial strains and growth conditions

All mutant strains constructed in this study are derived from *B. subtilis* 168 (*trpC2). Escherichia coli* MC1061 was used as intermediate cloning host for all plasmid constructions. Both *B. subtilis* and *E. coli* were grown aerobically at 37 °C in Lysogeny Broth (LB) unless otherwise indicated. When necessary, antibiotics were added to the growth medium as described previously (Cao et al., 2017).

### 2.2. Recombinant DNA techniques

Procedures for PCR, DNA purification, restriction, ligation and genetic transformation of *E. coli* and *B. subtilis* were carried out as described before (Konkol et al., 2013; Sambrook et al., 1989). Pfu X7 DNA polymerase (Norholm, 2010) was a kind gift from Bert Poolman, and the USER enzyme was purchased from New England Biolabs. All FastDigest restriction enzymes, Phusion and Dreamtaq DNA polymerases were acquired from Thermo Fisher Scientific (Landsmeer, Netherlands). The NucleoSpin Plasmid EasyPure and NucleoSpin Gel and PCR Clean-up kits were purchased from BIOKE (Leiden, Netherlands). All the reagents used were bought from Sigma unless otherwise indicated. Oligonucleotides were synthesized by Biolegio (Nijmegen, Netherlands). Sequencing of all our constructs was performed at MacroGen (Amsterdam, Netherlands).

### 2.3. Construction of strains and transcription factor mutagenesis libraries

A list of the plasmids and strains included in this work is presented in **Table 1**, and all the oligonucleotide primers used in this study are listed in **Table S1**. To create a tunable overexpression system, we introduced an expression cassette containing the β-galactosidase encoding gene (*lacZ*) into an IPTG inducible system (Claessen et al., 2008). This construct was integrated into the *mdr* locus by double homologous recombination, thus ensuring that only one copy of the enzyme-encoding DNA is present in the chromosome of *B. subtilis* 168. This reporter strain was termed JV156 (168_β-gal). All *B. subtilis* mutant candidates were derived from this reporter strain.

**Table 1.** Strains and plasmids used in this study.

| Strain or plasmid | Genotype | Reference or source |
| --- | --- | --- |
| *B. subtilis* |  |  |
| 168 | *trpC2* | (Kunst et al., 1997) |
| 168$_{\beta}$-gal | *trpC2, mdr::(P<sub>hyspank</sub>-lacZ spc<sup>r</sup>)* | This study |
| CodY<sup>-</sup> | *trpC2, codY::cm<sup>r</sup>, mdr::(P<sub>hyspank</sub>-lacZ spc<sup>r</sup>)* | This study |
| CcpA<sup>-</sup> | *trpC2, ccpA::erm<sup>r</sup>, mdr::(P<sub>hyspank</sub>-lacZ spc<sup>r</sup>)* | This study |
| CcpA<sup>-</sup>CodY<sup>-</sup> | *trpC2, ccpA::erm<sup>r</sup>, codY::cm<sup>r</sup>, mdr::(P<sub>hyspank</sub>-lacZ spc<sup>r</sup>)* | This study |
| CodY<sup>R214C</sup> | *trpC2, codY<sup>R214C</sup> cm<sup>r</sup>, mdr::(P<sub>hyspank</sub>-lacZ spc<sup>r</sup>)* | This study |
| CodY<sup>(E58A,I162V,R209H)</sup> | *trpC2, codY<sup>(E58A,I162V,R209H)</sup> cm<sup>r</sup>, mdr::(P<sub>hyspank</sub>-lacZ spc<sup>r</sup>)* | This study |
| CodY<sup>R214C</sup> CcpA<sup>-</sup> | *trpC2, ccpA::erm<sup>r</sup>, codY<sup>R214C</sup> cm<sup>r</sup>, mdr::(P<sub>hyspank</sub>-lacZ spc<sup>r</sup>)* | This study |
| CcpA<sup>T19S</sup> | *trpC2, ccpA<sup>T19S</sup> km<sup>r</sup>, mdr::(P<sub>hyspank</sub>-lacZ spc<sup>r</sup>)* | This study |
| CodY<sup>R214C</sup> CcpA<sup>T19S</sup> | *trpC2, ccpA<sup>T19S</sup> km<sup>r</sup>, codY<sup>R214C</sup> cm<sup>r</sup>, mdr::(P<sub>hyspank</sub>-lacZ spc<sup>r</sup>)* | This study |
| *E. coli* |  |  |
| MC1061 | F<sup>-</sup>, *araD139, Δ(ara-leu)7696, Δ(lac)X74, galU, galK, hsdR2, mcrA, mcrB1, rspL* | (Casadaban and Cohen, 1980) |
| BL21(DE3) | F<sup>-</sup>, *ompT, hsdS<sub>B</sub> (r<sub>B</sub><sup>-</sup>, m<sub>B</sub><sup>-</sup>), gal, dcm (DE3)* | Laboratory stock |
| Plasmids |  |  |
| pJV112 | *bla, mdr::(P<sub>hyspank</sub>-lacZ spc<sup>r</sup>), lacI* | This study |
| pUC18 | *bla, lacZ, amp<sup>r</sup>* | (Vieira and Messing, 1991) |
| pCH3 | pUC18_aroA_ccpA<sub>km<sup>r</sup></sub>_ytxD | This study |
| pCH3_CcpAKO | pUC18_aroA_erm<sup>r</sup>_ytxD | This study |
| pJV54 | pUC18_clpY_codY<sub>cm<sup>r</sup></sub>_flgB | This study |
| pJV55 | pUC18_clpY<sub>cm<sup>r</sup></sub>_flgB | This study |
| pDOW01 | amyE::(*P*<sub>hyspank</sub> spc<sup>r</sup>), lacI | (Weme et al., 2015a) |
| pDOW-CcpA | amyE::(*P*<sub>hyspank</sub>-ccpA<sub>his8</sub> spc<sup>r</sup>), lacI | This study |
| pDOW-CcpA<sup>T19S</sup> | amyE::(*P*<sub>hyspank</sub>-ccpA<sup>T19S</sup><sub>his8</sub> spc<sup>r</sup>), lacI | This study |
| pDOW-CodY | amyE::(*P*<sub>hyspank</sub>-his6codY spc<sup>r</sup>), lacI | This study |
| pDOW-CodY<sup>R214C</sup> | amyE::(*P*<sub>hyspank</sub>-his6codY<sup>R214C</sup> spc<sup>r</sup>), lacI | This study |

The mutant libraries of *codY* and *ccpA* were established successively as exemplified for *codY* in **Fig. S1B**. First, the integration vector was constructed with the USER (Uracil-Specific Excision Reagent) cloning method (Bitinaite et al., 2007). The obtained plasmid (pJV54), which consists of ~1 kbp flanking regions, antibiotic resistance marker and the pUC18 backbone, was applied as the template for mutagenesis experiments. The GeneMorph II Random Mutagenesis Kit (Agilent Technologies, United States) was utilized to achieve the desired mutation rate for each library according to the manufacturer’s instructions. Afterwards, the mutagenized *codY* sequences were cloned into pJV54, thereby replacing the native *codY* or *ccpA* gene and subsequently transformed into competent *E. coli*. Fresh transformants were pooled and the plasmids were extracted after overnight incubation. Finally, the plasmid DNA library was transformed into competent *B. subtilis* cells, and the mutagenized transcription factor genes were integrated into their native loci via double homologous recombination. In this way, the *codY\** and *ccpA\** libraries in *B. subtilis* were generated separately. The single and double knockout strains CodY^-^ (*codY::cm^r^*), CcpA^-^ (*ccpA::erm^r^*) and CcpA^-^CodY^-^ (*ccpA::erm^r^*, *codY::cm^r^*) were also constructed.

### 2.4. Black-white screening

All colonies of the *B. subtilis* mutant libraries were scraped off from the selective LB agar plates and collected into one flask with fresh LB medium supplemented with the appropriate antibiotic(s). After 37 °C overnight incubation, the culture was serially diluted and plated on selective agar medium consisting of Spizizen’s minimal medium (SMM) (Anagnostopoulos and Spizizen, 1961) with 1.0% glucose, 300 mg/l S-gal, 500 mg/l ferric ammonium citrate for black-white screening. After 20 h of incubation at 37 °C, colonies were isolated from the plates based on the color intensity and morphology, followed by sequence analysis and enzymatic assays.

### 2.5. β-Galactosidase assay

To determine the β-galactosidase activities, strains were grown in LB medium supplemented with 1.0% glucose and 0.1 mM IPTG, shaking at 37 °C and 220 rpm until the mid-exponential growth phase was reached (OD_600_ of 1.0). One milliliter of the cultures was harvested by centrifugation (14,000 rpm, 4 °C) and immediately frozen in liquid nitrogen. The pellet was processed for β-galactosidase quantitation as previously described (Smale, 2010). Each assay was performed in biological duplicates, and the mean value from three independent experiments was calculated.

### 2.6. Transcriptome analysis by RNA-Seq and qRT-PCR

Strains were grown in LB medium supplemented with 1.0% D-glucose and 0.1 mM IPTG until an OD_600_ of 1.0 was reached and 5 ml samples were harvested by centrifugation (7,000 rpm, 4 °C). The cell pellets were immediately shock frozen in liquid nitrogen. The total RNA was extracted as described earlier (Yi et al., 2017) and split into two aliquots for RNA sequencing and qRT-PCR. Sequencing of cDNA versions of the mRNAs was accomplished by PrimBio (USA), and the data analysis was performed as described before (de Jong et al., 2015; van der Meulen et al., 2016). For qRT-PCR, reverse transcription of the RNA samples was performed with the SuperScript^TM^ III Reverse Transcriptase kit, and quantitative PCR analysis was performed with the iQ5 Real-Time PCR Detection System (Bio-Rad) as described previously (Livak and Schmittgen, 2001).

### 2.7. Western blot analysis

Immunoblot analysis was performed according to previous studies (Brinsmade et al., 2014; Weme et al., 2015b). First, cells were grown to OD600 of 1.0, and 5 ml of the cultures were harvested by centrifugation (7,000 rpm, 5 min). The pellets were resuspended in 1 ml solution A [50 mM Tris·Cl (pH 7.5), 5% (vol/vol) glycerol, and 1 mM PMSF]. Subsequently, cells were broken by sonication for 2 min using 70% amplitude with 10-s bursts, 10-s pauses, and the total protein samples were collected by centrifugation (14,000 rpm, 5 min). Ten μg of total protein was heated for 5 min at 95 °C in denaturing loading buffer before being separated by SDS-PAGE, after which the proteins were transferred to a PVDF membrane (Millipore, USA). The membranes were blocked in PBST + 5% (wt/vol) BSA at 4 °C overnight. Subsequently, the membranes were separately subjected to a first incubation (90 min) with a rabbit anti-CodY (Ratnayake-Lecamwasam et al., 2001) and a rabbit anti-CcpA (Küster et al., 1996) polyclonal antibody (1:10,000) and a second incubation (90 min) with a donkey anti-rabbit IgG horseradish peroxidase (1:10,000) at room temperature. The signal intensity of bands was visualized using the ECL Prime kit (GE Healthcare Life Sciences) and detected by the Molecular Imager ChemiDoc XRS+ (Bio-Rad).

### 2.8. Electrophoretic mobility shift assay (EMSA)

*E. coli* BL21 (DE3) strains carrying the plasmid pDOW, in which different versions of the transcription factors were cloned, were grown until an OD_600_ of 0.7. After addition of the inducer IPTG (0.4 mM, final concentration), incubation was continued for six hours. Cells were harvested (4 °C, 7,000 rpm, 10 min) and then lysed with 1 mg/ml lysozyme and sonication. The His_6_-tagged CodY and CcpA proteins were purified using a Histrap^TM^ excel column following the manufacturer’s protocol (GE Healthcare Life Sciences). The purity of the transcription factor fraction was analyzed by sodium dodecyl sulfate (SDS)-polyacrylamide gel electrophoresis and Coomassie blue staining.

DNA probes were PCR amplified using Cy3-labeled primers, and the acquired PCR products were purified with the DNA clean-up kits (BIOKE). The DNA-target protein binding step was carried out in the presence of cofactors (2 mM FBP for CcpA, 2 mM GTP and 10 mM BCAAs for CodY) with 1X binding buffer, 0.2 μl of 10 mg/ml BSA, 5 nM of labeled DNA fragments and the purified His_6_-tagged proteins in different concentrations. The total volume was adjusted to 20 μl with MilliQ water and incubated at 30 °C for 20 min to complete the binding reaction. The obtained samples were loaded on a 5% nondenaturing polyacrylamide gel (750 μl 40% acrylamide, 600 μl 5× TBE, 5 μl TEMED, 50 μl 10% APS, MilliQ up to 6 ml). Electrophoresis was carried out in 0.5% TBE buffer (pH 7.4) at 200 V for 30 min. Afterward, fluorescence signals were recorded using a Fuji LAS-4000 imaging system.

## 3. Results

### 3.1. gTME libraries of CodY and CcpA allow selection of B. subtilis mutants with increased capacity of β-galactosidase production

The master transcriptional regulator CodY controls hundreds of genes in a large regulon, the products of which are mainly linked to nitrogen metabolism. More specifically, CodY senses the intracellular levels of BCAAs and GTP and represses or activates the transcription of nitrogen metabolic network related genes to trigger varying metabolic effects by binding to consensus function sites called the CodY box (Sonenshein, 2007). Therefore, any alteration of the CodY amino acid sequence can potentially reprogram the downstream metabolic fluxes and thus influence the production of heterologous proteins. To create a tunable overexpression system, we introduced an expression cassette containing the heterologous β-galactosidase encoding gene (*lacZ*) into an IPTG inducible system. The resulting construct P*_hyspank_-lacZ* was chromosomally integrated into the *mdr* locus of *B. subtilis* 168 to obtain the reporter strain *B. subtilis* 168_β-gal (**Fig. S1A**). Subsequently, by using a set of two error-prone DNA polymerases, three pools of randomly mutated *codY* genes with different mutation frequencies (1-4.5, 4.5-9, 9-16 mutations/kb amplified target gene) were generated. The mutagenized versions of *codY* were introduced into the P*_hyspank_-lacZ* fused reporter strain by double homologous recombination, thereby replacing the wildtype (WT) *codY* gene. In total, three independent CodY mutant libraries with a size of more than 5,000 clones were established. From isolates with a widely varying protein production potential, protein overproducing candidates were selected based on an increase in β-galactosidase activity (**Fig. 1**).

**Fig. 1.**
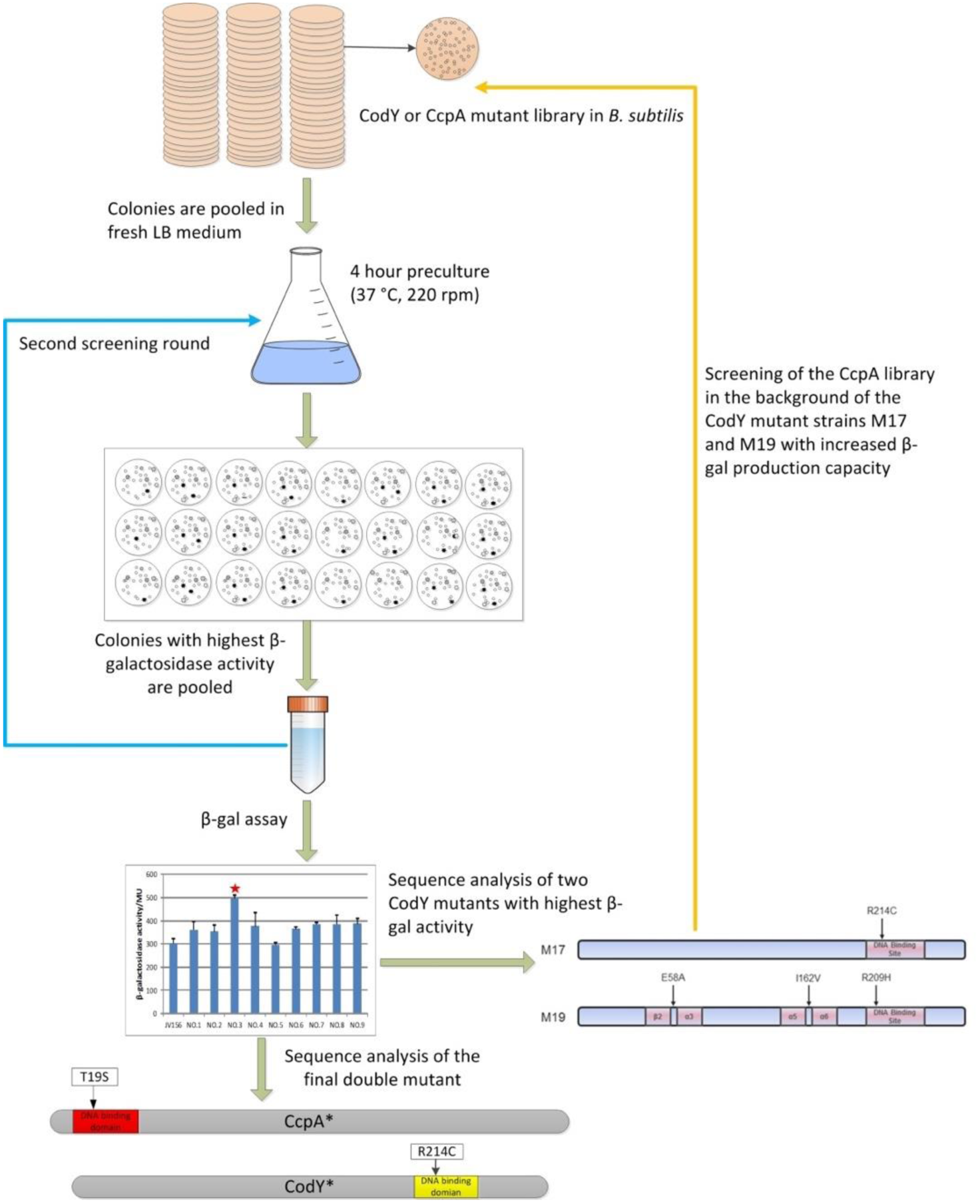
Workflow of the black-white screening for the identification of *B. subtilis* CodY or CcpA mutants with increased β-galactosidase production capacity. Desired phenotypes with a high conversion of the substrate S-gal that appeared as darkest colonies on the screening medium (SMM supplemented with 1.0% glucose, 300 mg/l S-gal, 500 mg/l ferric ammonium citrate) were selected and pooled. After a subsequent second screening round, the β-galactosidase activity of single mutants was quantitatively assessed in an enzymatic assay, and the mutants were analyzed by DNA sequencing. Finally, the targeted introduction of the CcpA^T19S^ mutation into the CodY^R214C^ mutant strain resulted in a strain of 290% enhanced the productivity of the heterologous enzyme.

The black-white screening was performed on transparent SMM plates, which allowed direct benchmarking of the color intensity and monitoring of the colony size. Phenotypes with increased β-galactosidase activity were selected out of a total of over 15,000 CodY mutant candidates, and the β-galactosidase production yields were quantified by colorimetric assays. *B. subtilis* mutants with higher reporter enzyme production originated from the low mutation frequency library (1-4.5 mutations/kb), indicating that libraries with more than 4.5 mutations/kb have a higher probability of harboring phenotypic variations that negatively affect the biosynthesis of the reporter protein. Finally, two CodY mutants showing the highest improvement in production capacity, i.e. M17 and M19, were found to generate 52% and 40% more β-gal than the reporter strain equipped with the unmodified, wildtype *codY* gene (**Fig. S2A**). In both mutants, we identified one amino acid substitution within the DNA-binding HTH motif of CodY. In M17, solely the single mutation R214C was detected, whereas M19 carried the HTH domain mutation R209H next to the amino acid exchanges E58A and I162V (**Fig. S2B**). Hence, we showed that the gTME approach coupled to an enzymatic protein production capacity screen could be successfully applied in *B. subtilis* and that specific mutations within the DNA binding domain of CodY resulted in a significantly increased product yield of the heterologous protein β-galactosidase.

To explore whether the production potential of the previously engineered expression hosts could be further improved, we additionally reprogrammed the carbon metabolic network in the two CodY mutants by random mutagenesis of the transcriptional regulator CcpA. A *ccpA*\* library with 1-4.5 mutations/kb mutation frequency was constructed and integrated into the chromosome of M17 and M19 to obtain two libraries, M17 (*ccpA*\*) and M19 (*ccpA*\*). Subsequently, CcpA mutant strains were selected from each library by the black-white screening based on their higher β-galactosidase product yields. The two best performing phenotypes from the two different libraries showed the same amino acid exchange, T19S, in the DNA-binding HTH motif of CcpA (**Fig. S3**). To further verify the influence of the single amino acid substitution T19S and to rule out the possibility of acquisition of additional (compensatory) mutations, we performed site-directed mutagenesis of *ccpA* to introduce the mutation T19S in different background strains, thereby obtaining the mutants CcpA^T19S^ and M17CcpA^T19S^. Since then, the M17 and M17CcpA^T19S^ were renamed as CodY^R214C^ and CodY^R214C^CcpA^T19S^ in the following analyses. In comparison to the parental WT control strain 168_β-gal, all transcription factor mutants showed significantly increased β-gal activities (**Fig. 2**). We observed that the deficiency of CcpA and/or CodY, which was introduced by gene knockouts, could already improve the β-galactosidase yield by 10-20% than that of WT strain. However, in the single mutants CodY^R214C^ and CcpA^T19S^, the respective production of the target enzyme was increased to 152% and 140% relative to the WT. This indicates that single mutations in the DNA-interacting HTH domains of the global transcriptional regulators, rather than complete gene knockouts, were advantageous. Moreover, these production advantages were synergistic when the *codY* and *ccpA* mutations were combined in one strain, because heterologous protein production was enhanced to 290% in the double mutant CodY^R214C^CcpA^T19S^. In line with the results presented above, a significant increase of GFP (89%), XynA (xylanase protein from *B. subtilis*) (more than 11-fold) and PepP (aminopeptidase P from *Lactococcus lactis*) (80%) production could be achieved in CodY^R214C^CcpA^T19S^ relative to the WT host (**Fig. S4**). This demonstrates that productivity improvement in the genetically modified cell factory is achieved for various proteins.

**Fig. 2.**
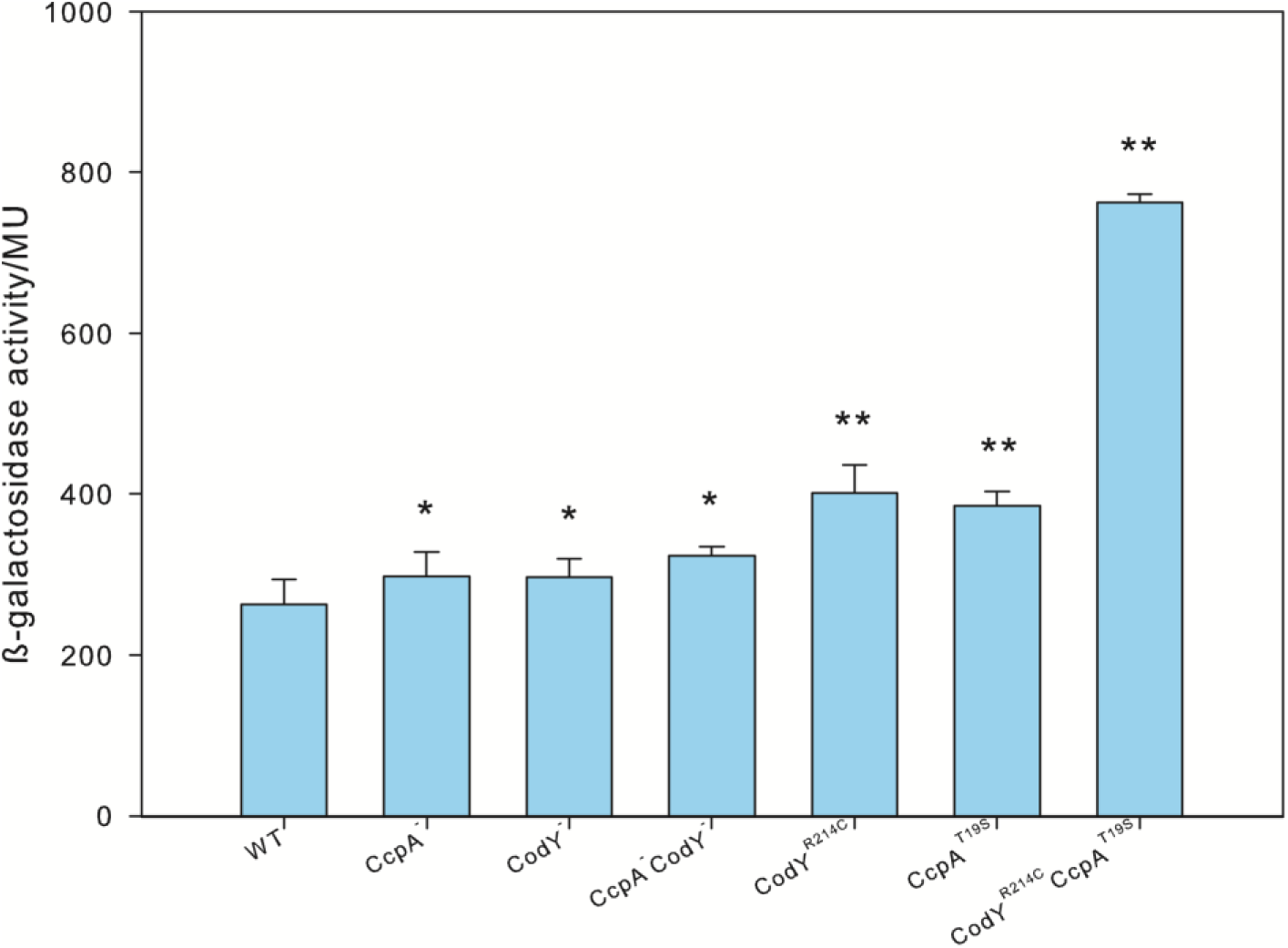
β-galactosidase activities in *B. subtilis* strains with null mutations or single amino acid substitutions in the DNA-binding HTH domain of CodY and CcpA. Enzymatic assay of the recombinant strains was carried out in comparison to the control strain carrying a *lacZ* gene as a capacity monitor (WT). The cultures of were sampled at an OD_600_ of 1.0, and the β-galactosidase activities are shown in Miller Units. Each column represents the mean ± SD of three independent experiments. The statistical significance of differences was performed by the T-TEST, the black symbols represent the comparison with the WT (*P<0.05, **P<0.01).

### 3.2. Enhanced transcription of specific operons involved in nitrogen metabolism is positively correlated with β-galactosidase production

The target genes that are under the regulation of CodY and CcpA have been identified by genome-wide analyses of the transcriptome and protein-DNA interactions in *B. subtilis* in various studies (Belitsky and Sonenshein, 2008, 2013;Blencke et al., 2003; Brinsmade et al., 2014; Geiger and Wolz, 2014; Marciniak et al., 2012; Molle et al., 2003; Moreno et al., 2001). We opted to investigate the global cellular response to the amino acid substitutions in the DNA-binding HTH motifs of the transcriptional regulators CodY and CcpA during the mid-exponential growth phase to obtain a better understanding of the regulatory effects. The transcriptome patterns associated with the selected strains implied that the expression of individual genes is differentially affected, which would display a broad range of sensitivities to the HTH sequence mutations (**Fig. S5**). This would be expected because of the scale of global transcriptional regulation and the complexity of the interplay between diverse metabolic networks (Büscher et al., 2012) and also because CodY binds to different target sites with a varying affinity (Brinsmade et al., 2014). Next to genes that are known regulon members and under direct transcriptional control of CodY or CcpA, additional genes involved in nitrogen and carbon core metabolic pathways were transcribed differentially in the mutant strains. These were clustered according to their functional category in the *Subti*Wiki database (**Fig. S6**).

In comparison to the WT strain, the vast majority of CodY regulon members and nitrogen metabolism associated genes were either up-regulated or unchanged in the strains with mutations in DNA-binding regions of CodY and/or CcpA (**Fig. S6A-D**). In contrast, the overall fluctuation in expression levels of genes from the central carbon metabolism was modest; most of the genes were slightly down-regulated, and only a few were expressed 2-fold higher than in the WT (**Fig. S6E**). Interestingly, a specific set of gene clusters was positively correlated with the β-galactosidase production performance in the HTH domain mutation strains (**Fig. 3**). All of these operons are negatively controlled by CodY, and their corresponding products are involved in the uptake and utilization of specific nitrogen sources. The operons *rocABC* and *rocDEF* encode enzymes that participate in the uptake and utilization of arginine, ornithine, and citrulline (Calogero et al., 1994; Gardan et al., 1995). The *hutPHUIGM* operon is involved in histidine metabolism, which is additionally also negatively regulated by CcpA (Gopinath et al., 2008). The *appDFABC*operon is directly related to nitrogen source utilization by encoding an ABC transporter for the uptake of peptides (Koide and Hoch, 1994). As illustrated in **Fig. 3,** CodY^R214C^ and CcpA^T19S^ separately promoted the alteration of specific pathways for nutrient uptake and utilization, and these benefits were clearly synergistic in the double mutant host. Thus, we demonstrate that mutations in the conserved DNA binding motifs of CodY and CcpA significantly enhanced the expression of specific operons, and the resulting up-regulated nitrogen source metabolism was identified as a beneficial factor allowing the mutant strains to synthesize more reporter protein during growth in LB supplemented with 1.0% glucose.

**Fig. 3.**
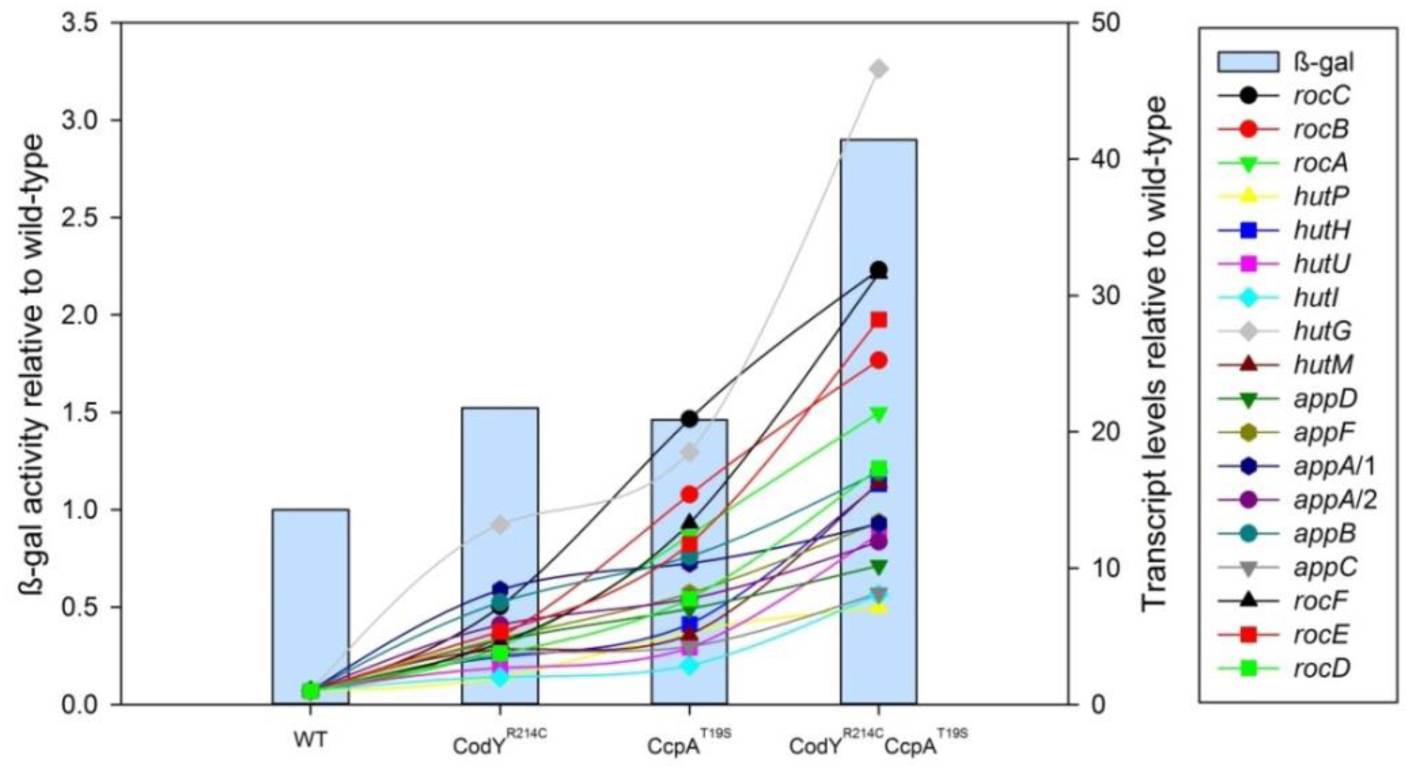
Transcription levels of the operons *rocABC*, *hutPHUIGM*, *appDFABC* and *rocDEF* are correlated with increased β-galactosidase production. The enzymatic activity of β-galactosidase is shown relative to the enzyme activity in the WT (blue columns). Because of the high differences in transcript levels, the normalized RPKM (Reads Per Kilobase per Million mapped reads) values of indicated genes are shown relative to the WT levels, which were arbitrarily scaled to 1.

### 3.3. Stronger repression of carbon metabolic pathways and de-repression of nitrogen metabolic pathways benefit the synthesis of β-galactosidase

Since the HTH motifs of CodY and CcpA are highly conserved among many low G+C Gram-positive species (Joseph et al., 2005; Pabo and Sauer, 1992), we next addressed the question whether the T19S and R214C mutations affect the DNA-binding ability of the regulator proteins. Gel electrophoresis mobility shift assays (EMSAs) with purified CcpA and CodY WT and mutant proteins and *ackA* and *ilvB* promoter fragments revealed that all protein variants were capable of binding to the selected DNA probes (**Fig. 4A**), which was in line with previous findings (Shivers et al., 2006; Shivers and Sonenshein, 2005). The CodY^R214C^ and CcpA^T19S^ mutants, however, bound to several regulatory sites with reduced and increased efficiency respectively, in comparison to their WT proteins. Furthermore, the mutation T19S significantly enhanced the binding efficiency of CcpA to the promoter regions of *rbsR* and *treP,* which are under the direct negative control of CcpA (**Fig. S7**).

The transcriptome analysis further revealed that the transcription factors CodY and CcpA themselves were differentially expressed in the reprogrammed HTH domain mutants. Surprisingly, these two regulatory proteins displayed exactly opposite expression patterns in the strains (**Fig. 4B**). The transcript abundance of CodY was around three times higher than that of CcpA in the WT strain JV156, and this difference in transcript levels was halved in the CodY^R241C^ mutant. Particularly, the CcpA mutation T19S led to an increase in transcription of *ccpA* and substantially decreased the transcription of *codY*, making the CcpA^T19S^ mutant strain nearly behave in the direction of a CodY deficient strain. However, the differential accumulation of the two transcriptional regulators got balanced when these two mutations were combined in one cell (**Fig. 4B**). This is supposedly a consequence of the interplay between these two regulators with altered regulation efficiencies and expression capacities. Importantly, this observation could be confirmed at the transcriptional and translational level by qRT-PCR and Western Blotting (**Fig. 4C** and **Fig. 4D)**.

**Fig. 4.**
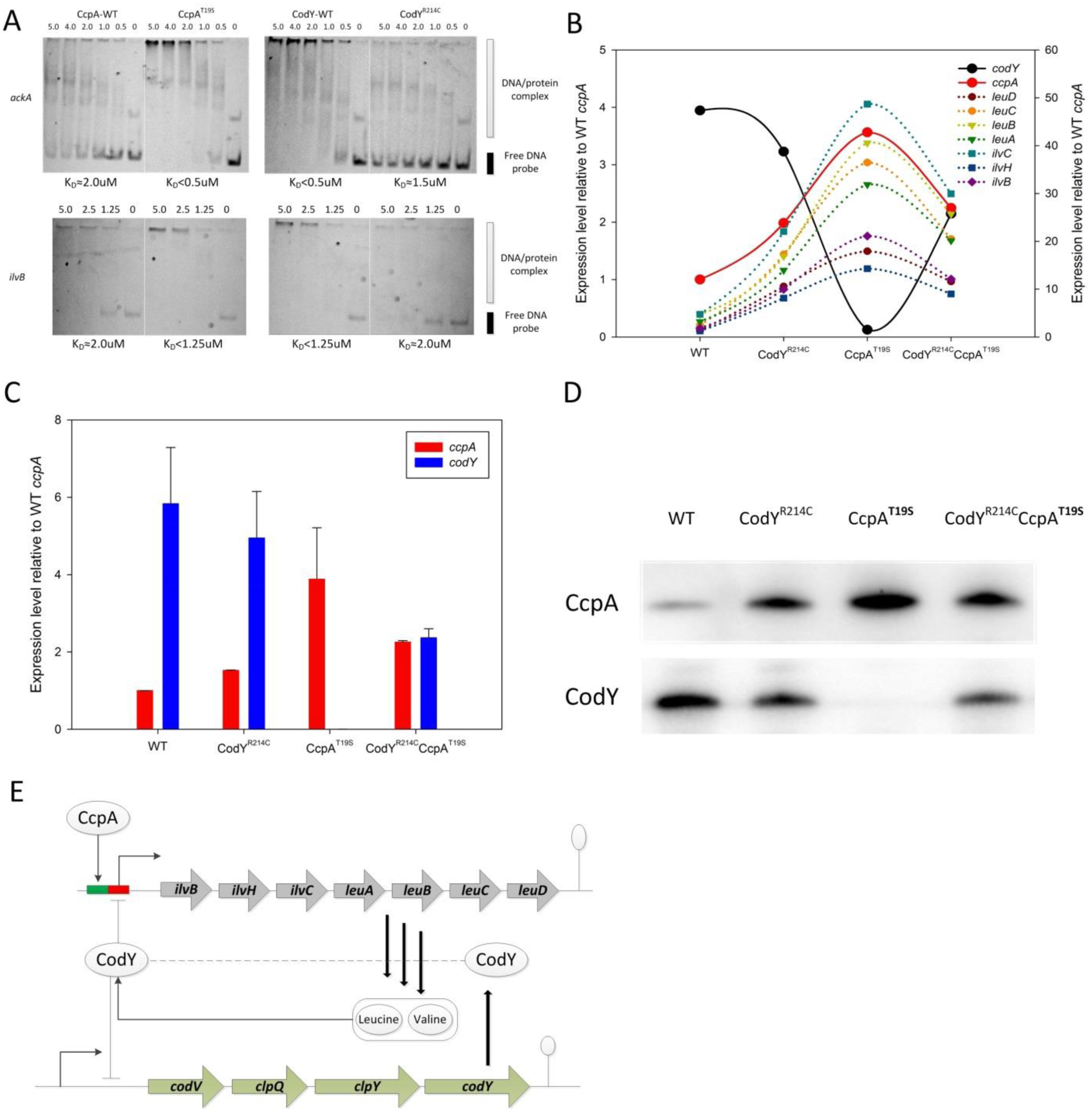
(A) Gel mobility shift analysis of CodY and CcpA binding to the regulatory regions of *ackA* and *ilvB*. The 3’ ends of DNA fragments were labeled with Cy3, and the obtained DNA probes (0.1 μM) were incubated with various concentrations (μM) of His_6_-tagged transcriptional factors. K_D_ reflects the protein concentration needed to shift 50% of DNA fragments (Belitsky, 2011). (B) The relative expression levels of CodY, CcpA and the *ilv-leu* operon in the HTH domain mutant strains. The RPKM values of each strain were normalized by that of *sigA* (internal reference gene). The normalized values of *ccpA*, *codY* and the *ilv-leu* operon genes were related to WT *ccpA*, and the value of WT *ccpA* was finally defined to 1. (C) qRT-PCR to analyze the expression level of the two proteins. The transcript level of *sigA* was used as the internal control, and the values obtained were related to that of WT *ccpA*, which was normalized to 1.0. Each column represents the mean ± SD of three independent experiments, and each assay was performed in duplicate. (D) Immunoblots of lysates from cells grown to an OD_600_ of 1.0. Strains were grown in LB medium in the presence of 1.0% glucose and 0.1 mM IPTG; the proteins were separately immunoblotted with polyclonal CcpA and CodY antibodies. (E) Schematic diagram of the interaction between CcpA and CodY mediated by the biosynthesis of BCAAs. Arrows and perpendiculars represent the positive and negative actions, respectively.

We thus show that the bacteria tend to alter the overall metabolic network fluxes through the expression variation of these two regulators to meet the demand of resources for the overproduction of heterologous proteins. In other words, the metabolic shifts occurring in *B. subtilis* can be regarded as a fitness adaptation of this microbial cell factory to the global transcriptome perturbations and the requirement of high-yield protein production (Goel et al., 2012). Finally, the increased CcpA and decreased CodY protein levels lead to an enhancement of the repression of the carbon metabolism and amplification of the reactions in the nitrogen metabolism networks, and thus the reprogrammed metabolic networks are obviously beneficial for the biosynthesis of β-galactosidase.

### 3.4. CcpA^T19S^ promotes β-galactosidase synthesis by impacting co-factor availability and the auto-regulatory expression loop of CodY

In *B. subtilis*, the two BCAAs leucine and valine, which are biosynthesized through the catalysis of *ilv-leu* encoded enzymes, effectively increase the affinity of CodY to its DNA binding sites. CodY and CcpA bind to different regulatory regions of the *ilv-leu* gene cluster promoter region and CcpA thus indirectly controls the expression of other CodY-regulated genes by the modulation of the intracellular level of BCAAs (Shivers and Sonenshein, 2005; Tojo et al., 2005). The transcription profile of the HTH mutants reflects that CcpA and CodY regulated metabolic pathways interact at the node of the *ilv-leu* operon (**Fig. 4B**). The two opposite transcriptional regulatory effects cause this operon to be expressed in a similar pattern as the *ccpA* gene but exactly opposite to *codY* (**Fig. 4B**). More specifically, the higher levels of accumulated BCAAs have negative feedback on the expression of CodY itself (**Fig. 4E**). The increased β-galactosidase production in the CodY^R214C^ and/or CcpA^T19S^ strains was strongly correlated with the transcript abundance of several CodY-regulated operons, which is obviously caused by an indirect derepression mediated by CcpA^T19S^ (through indirectly suppressing the transcription and activity of CodY^R214C^ (**Fig. S8)**). Hence, these two global regulatory proteins with altered binding specificities and expression activities cooperate to achieve the synergistic effect of target protein expression in the double mutant strain CodY^R214C^CcpA^T19S^ by prioritizing the upregulation of specific CodY targets via the modulation of BCAAs biosynthesis.

### 3.5. The gene regulatory network of nitrogen metabolism is more affected in high-capacity production mutants than the network of carbon metabolism

A higher number of CodY-regulated genes than CcpA regulon member genes were altered with respect to their expression levels in various mutants relative to the WT strain (**Fig. S6A and Fig. S6B)**. Next to the *ilv-leu* operon, 52 CodY-repressed genes including the BCAA biosynthesis related genes *ilvA, ilvD, and ybgE* were differentially expressed (Fink, 1993; Shivers and Sonenshein, 2005), while only two CcpA targets (*sacA* and *sacP*) showed a consistent co-expression with CcpA (**Fig. S9**). We therefore conclude that CodY-regulated genes react more sensitively to the changes of the corresponding regulator levels, while the expression levels of CcpA regulon genes remained rather stable in all HTH domain mutant strains. However, by specifically analyzing genes related to amino acid utilization or biosynthesis/acquisition and carbon core metabolism according to the gene ontology (GO) classification in *Subti*Wiki database, significantly in **Fig. S6C**, **Fig. S6D** and **Fig. S6E**, more amino acid metabolism genes were differentially expressed than carbon core metabolic genes. In short, the response of amino acid metabolic pathways to the changed transcriptome is more significant than that of central carbon metabolism in overproduction strains.

## 4. Discussion

Decades of research have demonstrated the importance of improving the cell factory protein productivity by specific target modification, but less work has been undertaken on engineering the global transcription machinery of central metabolic networks. Such gTME strategy, which focuses on the increase of end-products by perturbing the global transcriptome and rerouting metabolic fluxes at a global level, can remarkably simplify the enhancement strategy even without a thorough understanding of the underlying metabolic regulatory mechanisms (Lanza and Alper, 2012; Woolston et al., 2013). In this study, we developed the toolkit of gTME and high-throughput screening, which proved to outperform the traditional approaches for increasing the production of microbial cell factories. Theoretically, this tailor-made system offers potential to overexpress any heterologous protein by achieving multiple and simultaneous perturbations of the whole transcriptome and metabolome. Next to the model protein β-galactosidase, we also managed to overproduce additional heterologous proteins (GFP, xylanase, and peptidase) in the background of the transcription factor mutant. The cell factory productivity can be further influenced by the intracellular nutrient availability, codon usage and the utilization bias for some specific nitrogen sources. Although the level of gTME-based strain improvement differs per protein used, this strategy illustrates the broader application scope in overproducing a variety of proteins.

Most importantly, the system-wide analyses involving transcriptomics and protein-DNA binding assays provide a better understanding of the complex interactions between central metabolic pathways in *B. subtilis*. In contrast to the nitrogen metabolic network, the central carbon metabolism was less responsive to the transcriptional regulatory perturbations in all the mutants. This reflects that the threshold activities of regulators required for each gene are unequal, and the individual targets for both regulators are subject to the differential, gradual stimulation and repression (Brinsmade et al., 2014). Our data suggest that the higher activity of CcpA^T19S^ exceeded the maximum activation threshold, which was obviously sufficient to keep a stringent inhibition of gene expression of a vast majority of the CcpA regulon. In contrast, the activity of CodY^R214C^ was decreased and obviously lower than the level that full repression of gene transcription demands. Consequently, a set of CodY targets was dramatically upregulated due to the decline or even elimination of the transcriptional and metabolic repression. Although these two transcription factors regulate more than 200 genes directly (Michna et al., 2015), only one out of ten genes showed an altered expression profile in response to the global transcriptome perturbations. This is likely based on the fact that the vast majority of genes and operons are subject to complex, multiple forms of regulation at different expression levels.

In the natural environment, the availability of nutrients can be highly variable and the bacteria have evolved sophisticated adaptation systems for making good use of a wide range of sources of essential elements (Sonenshein, 2007). Therefore, the carbon core metabolism, which can guarantee essential energy and building blocks supply (Chubukov et al., 2014), has been well evolved to serve the bacteria in various conditions. The central pathways can be protected against the stochastic fluctuations by the overabundance of relevant enzymes (Kochanowski et al., 2013). The generation of buffer space ensures that the transcriptome perturbations will not severely restrict the capabilities of cellular energy metabolism. During the long-lasting natural evolutionary process, surviving under unfavorable or extreme growth environments is the primary task for microorganisms in contrast to the human demand for overproduction of heterologous protein, explaining why a global adjustment of global N- and C-metabolism is effective to support the latter.

In brief, we could significantly improve the productivity of *B. subtilis* by the rewiring of central metabolic regulation, which promotes a good balance of resource distributions between normal cellular processes and needs for heterologous protein production. Undoubtedly, further improvements in our ability to reveal the underlying interactions between transcriptional regulation and dynamic metabolic status will come from future studies (Kochanowski et al., 2013). This investigation provides a new approach to improve *B. subtilis* as a cell factory, which is of broad significance for both industrial application and fundamental studies.

## Acknowledgments

We thank Tjeerd van Rij (DSM) and Marc Kolkman (Genencor) for helpful discussions. This research was partially funded by a grant from the former Kluyver Center for Genomics of Industrial Fermentation (Delft/Groningen) to JVH and by an ALW grant to RDOW. HC was supported by a grant from China Scholarship Council (CSC). We thank Dr. Lance Keller (Department of Fundamental Microbiology, University of Lausanne) for proofreading the manuscript. We are grateful to Anne de Jong (Department of Molecular Genetics, University of Groningen) for expert technical assistance of transcriptome analysis.

